# CoRINs: A tool to compare residue interaction networks from homologous proteins and conformers

**DOI:** 10.1101/2020.06.29.178541

**Authors:** Felipe V. da Fonseca, Romildo O. Souza Júnior, Marília V. A. de Almeida, Thiago D. Soares, Diego A. A. Morais, Rodrigo J. S. Dalmolin, João Paulo M. S. Lima

## Abstract

**Motivation:** A useful approach to evaluate protein structure and quickly visualize crucial physicochemical interactions related to protein function is to construct Residue Interactions Networks (RINs). By using this application of graphs theory, the amino acid residues constitute the nodes, and the edges represent their interactions with other structural elements. Although several tools that construct RINs are available, many of them do not compare RINs from distinct protein structures. This comparison can give valuable insights into the understanding of conformational changes and the effects of amino acid substitutions in protein structure and function. With that in mind, we present CoRINs (Comparator of Residue Interaction Networks), a software tool that extensively compares RINs. The program has an accessible and user-friendly web interface, which summarizes the differences in several network parameters using interactive plots and tables. As a usage example of CoRINs, we compared RINs from conformers of two cancer-associated proteins.

**Availability:** The program is available at https://github.com/LasisUFRN/CoRINs.

## INTRODUCTION

The study of protein structure and its function relationships rely on the principle that protein tridimensional structure is dependent on its amino acid residues sequence and their interatomic interactions (Anfisen, 1973; Kuhlman and Bradley, 2019). These later exert crucial importance on the understanding of native conformations, since highly dissimilar protein sequences can give rise to the same fold (Vijayabaskar and Vishveshwara, 2012). These analyses usually require molecular modeling techniques and the visual inspection of multiple tridimensional structure files in molecular visualization programs. However, assessing the structural effects of sequence divergence between homologous proteins, polymorphisms, and mutations on the protein function through the comparison of multiple PDB files can be daunting, especially in automated pipelines.

A useful approach to overcome this and to rapidly evaluate protein structure is the construction of Residues Interaction Networks (RINs). Also referred to as protein structure networks (Serçinoglu and Ozbek, 2018), this approach applies graph theory to protein structure. In RINs, each node represents a residue, and their edges represent chemical and physical interactions with other elements of the structure (Martin et al., 2011). Therefore, using standard network parameters such as degree, betweenness, and clustering coefficient, one can rapidly deduce the importance of any given amino acid residue to the overall conformational structure of the protein or its specific function on activity or catalysis (Piovesan et al., 2016). Several available tools construct RIN from a protein structure file input, such as RING (Martin et al., 2011), RING2.0 (Piovesan et al., 2016), RINalyzer (Doncheva et al. 2011), ProSNEX (Aydinkal et al., 2019), and others that can couple these analyses to several conformational variants calculated from molecular dynamics experiments (MD), such as gRINN (Serçinoglu and Ozbek, 2018) and RIP-MD (Contreras-Riquelme et al., 2018).

Here, we present a new tool, CoRINs (Comparator of Residue Interaction Networks), which compares RINs from different protein structures, and summarizes their differences on network formalism parameters for each residue. CoRINs perform a pairwise comparison of each input RIN and display the main differences between them. We believe that CoRINs reports are useful in the evaluation of protein conformational variation and as an additional tool to validate models from homology modeling.

## PROGRAM OVERVIEW

CoRINs requires as input RING2.0 (Piovesan et al., 2016) output files for the same protein in different conformations or from a set of PDB files from homologous proteins. Both text files, nodes.txt, and edges.txt, or the generated compressed files (.zip), can be used as input. The core development of CoRINs used the Pandas library due to its high performance in data analyses. For the tool’s backend and front-end development, we chose the Python language using the frameworks Django (https://www.djangoproject.com) and Bootstrap (https://getbootstrap.com/), respectively. For graphics construction and overall layout, we used the D3.js (https://d3js.org/) JavaScript library.

We designed the CoRINs interface focusing on users that need to evaluate the differences from protein’s RINs rapidly, and are not familiar with or do not want to use scripting languages or other network-construction programs, like Cytoscape (Cline et al., 2007). From the CoRINs home screen, you can submit the RINs’ files using the “Load” button, which will open a file manager according to the operating system used. CoRINs then converts the input nodes and edges files in data frames, and pairwisely compares each protein chain according to the amino acid position. It also compares RINs from chains within the same protein. The tool lists all RINs’ differences in the output of five distinct text files. The reports include the differences between nodes, amino acid changes, differences between edges, a comparison of all degree values and their variation at each position, and additional network parameters, as clustering coefficient and betweenness weighted. The program calculates these two parameters using a custom script and R packages (See supplementary methods).

CoRINs’ interface also interactively displays the results, using searchable tables and summarizing plots (Figure 1). The first overview of the RINs comparative consists of plots representing the standard deviation of degrees’ values from each residue (Figure 1A). Two bar charts display the plotted overall differences in the number of chemical interactions and the ten more significant degree differences per residue from any pair of RINs (Figure 1C). Also, the user can have a quick and interactive overlook of the total RINs’ differences per residue and degree values in heat map-like plots (Figure 1D), with squares representing each protein sequence position and the parameter difference displayed on a color scale. A few other searchable tables display the different amino acids, the unique interactions, and the betweenness weighted and clustering coefficient (Figure 1B). These features are useful for conformers analyses, as described in the following use example, to the validation of homology structure models or to assess the impact of amino acid changes.

**Figure 1 -.**
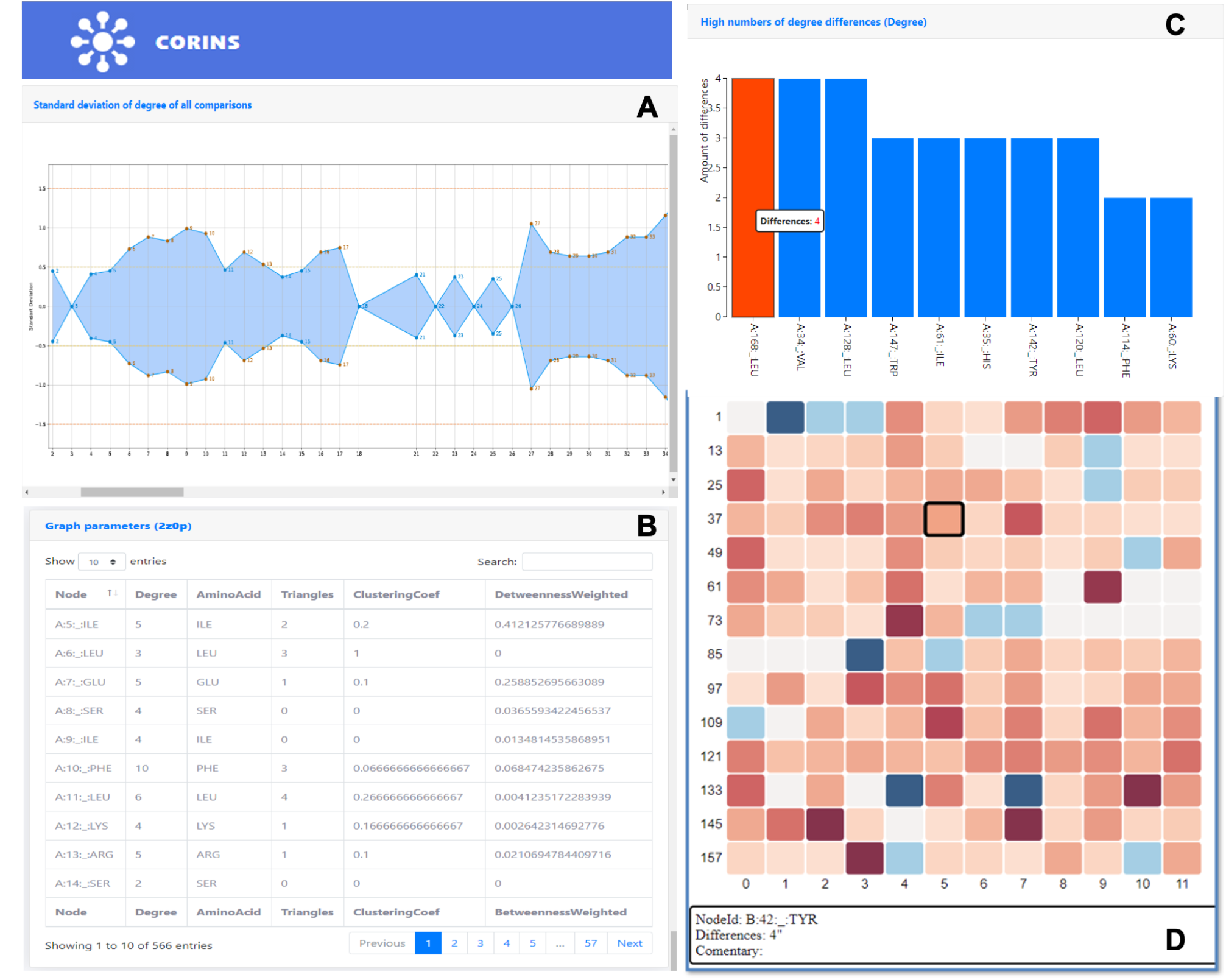
Screenshots from the CoRINs program interface. (A) Plot representing the standard deviation of degrees’ values from each residue from the comparison of all the submitted RINs. (B) One of CoRINs’ interactive tables, displaying additional network parameters. (C) Bar chart describing the more significant degree differences per residue from a pair of RINs. (D) Interactive plot displaying an overlook of the total RINs’ differences per residue.

## USAGE EXAMPLE

Using CoRINs, we analyzed several conformers from two cancer-associated genes listed in the CoDNaS database (Monzon et al., 2016). The protein BRAF (UniProtKB - P15056) is a cytosolic serine/threonine kinase of the rapidly accelerated fibrosarcoma (RAF) family that is involved in the transduction of mitogenic signals and associated to several human cancers (Dankner et al., 2018). We analyzed nine conformers from the BRAF kinase domain (pos. 457-717) clustered into the same pool (ID_POOL_CoDNaS 4MNF_A). The other chosen protein was the Phosphatidylinositol 3,4,5-trisphosphate 3-phosphatase (PTEN - UniProtKB - P60484), a dual-specificity phosphatase related to cell growth and survival (Chen et al., 2018). There is a single ensemble of thirteen conformers of PTEN, from 4 different PDB files (ID_POOL_CoDNaS 1DR5_A).

From CoRINs graphical display of the standard deviation of the residues’ degree values (Dsd), we assessed the protein sites that presented a higher variation in chemical interactions. This plot categorizes these values into four groups: Dsd=0, describing the residues that do not shown variation in degree values; 0 < Dsd ≤ 0.5 (blue color), describing the residues with low variation in chemical interactions; 0.5 < Dsd < 1.5 (brown color), describing residues with considerable changes in their connections; and Dsd ≥ 1.5 (red color), remarking probable positions of conformational variations. Then, to verify the variation in residue degree values in protein sites more prone to mutations, we retrieved missense mutations from The Cancer Genome Atlas (TCGA) and ClinVar (Landrum et al., 2016) databases.

Table 1 lists the number of mutations from both databases mapped into the three mentioned categories of variation. In BRAF, most missense mutations reported from TCGA occur in sites that present considerable variation in the number of non-covalent interactions. Moreover, five of the nine probable pathogenic mutations retrieved from ClinVar fell into residues positions with higher variation in connections. This same analysis, using the reported TCGA mutations of the PTEN conformers, showed similar results. However, for this protein, half of the mutations reported from ClinVar mapped to positions with a presumable little conformational variation.

**Table 1 -.** Number of mutations from TCGA and ClinVar databases versus the standard deviation of the residues’ degree values (Dsd).

| Dsd values | Mutation count - BRAF protein |  | Mutation count - PTEN protein |  |
| --- | --- | --- | --- | --- |
|  | TCGA | ClinVar | TCGA | ClinVar |
| **Dsd=0** | 1 | 0 | 12 | 0 |
| **$0 < \text{Dsd} \leq 0.5$** | 3 | 0 | 52 | 5 |
| **$0.5 < \text{Dsd} < 1.5$** | 27 | 4 | 131 | 2 |
| **$\text{Dsd} \geq 1.5$** | 35 | 5 | 53 | 3 |

## CONCLUSION

In conclusion, CoRINs comparative analyses from protein conformers can help identify critical physicochemical interactions to protein function. Also, coupled with homology modeling, the tool’s results can be useful to assess the putative impacts of a mutation on the protein structure and activity, by identifying its effects on the protein network.

## ACKNOWLEDGMENTS

The authors are indebted to the High-Performance Computing Center (NPAD) at UFRN for the availability of computational resources and the Instituto Metropole Digital (IMD) for the support in the realization of this work. The analysis demonstrated here used data generated by the TCGA Research Network: https://www.cancer.gov/tcga.

## AUTHOR CONTRIBUTIONS

FV Fonseca and JPMS Lima conceptualized the program. FV Fonseca was responsible for CoRINs’ core development. FV Fonseca and TD Soares elaborated the web design. RO Souza Júnior revised the main code and scripts and containerized the application. FV Fonseca and MVA Almeida worked in the protein analyses. DAA Morais and RJS Dalmolin suggested the additional network parameters and wrote R scripts. FV Fonseca and JPMS Lima wrote the paper.

