## Supplementary Methods for "CoRINs: A tool to compare residue interaction networks from homologous proteins and conformers"

**Availability:** The program is available at: https://github.com/LasisUFRN/CoRINs.

### SUPPLEMENTARY METHODS

To construct the Residue Interaction Networks (RINs), you need to use the RING2.0 program or webserver (Piovesan et al., 2016) available at http://old.protein.bio.unipd.it/ring/. For now, CoRINs only uses RING2.0 output files as input, but we pretend to expand this option soon.

To calculate the clustering coefficient and betweenness weighted parameters for each node, CoRINs uses RING2.0 generated file edges.txt. To calculate the clustering coefficient, we use the following equation:

$$\frac{Number\ of\ triangles\ of\ node_{(i)}}{0.5 \times Degree\ of\ node_{(i)} \times (Degree\ of\ node_{(i)} - 1)} \quad (1)$$

CoRINs program then counts the number of triangles using the function count_triangles of the R package igraph (Csardi and Nepusz, 2006 - http://igraph.org/). The estimated clustering coefficient ranges from 0 to 1, where 1 is the maximum possible number of connections. In CoRINs, we normalized the betweenness centrality utilizing the distance between each amino acid (edges) as weight and calculated it using the betweenness function of igraph (Csardi and Nepusz, 2006 - http://igraph.org/). The results range from 0 to 1. Therefore, a value of 0.1 means that 10% of the shortest pathways of the network pass through that node.

For the usage example, we used RING-constructed RINs from the following PDB files, listed as conformers at the CoDNas database (Monzon et al., 2016):

- BRAF protein (ID_POOL_CoDNaS 4MNF_A): 3OG7, 4CQE, 4XV1, 4XV2, 4XV9, 5HID, 5ITA, 5JRQ, 5JSM.
- PTEN protein (ID_POOL_CoDNaS 1DR5_A): thirteen conformers from the PDBs 1D5R, 5BUG, 5BZX, 5BZZ.
